# Efficient delivery of small RNAs to podocytes *in vitro* by direct exosome transfection

**DOI:** 10.1101/2024.10.11.617856

**Authors:** Tim Lange, Luzia Maron, Claudia Weber, Rabea Schlüter, Nicole Endlich

**Affiliations:** Institute of Anatomy and Cell Biology, University Medicine Greifswald, Greifswald, Germany; Imaging Center of the Department of Biology, Greifswald University, Greifswald, Germany

## Abstract

Podocytes are a crucial component of the glomerular filtration barrier and changes in their 3D structure contribute to over 80% of chronic kidney disease (CKD) cases. Exosomal small RNAs play a key role in cell-cell communication in CKD and may serve as nanocarriers for delivering small RNAs into podocytes. However, the uptake of exosomal cargo by podocytes remains poorly understood. This study explores the use of isolated exosomes, directly transfected with fluorescently labeled small RNAs, for tracking and delivering small RNAs to cultured podocytes.

Exosomes were isolated from immortalized murine podocytes and transfected with Cy3-labeled siRNA and miRNA controls using the ExoFect siRNA/miRNA Transfection Kit. We characterized the transfected exosomes via transmission electron microscopy (TEM) and Western blot for exosomal markers CD9 and TSG101. Subsequently, we co-cultured these exosomes with podocytes and used confocal laser-scanning microscopy (cLSM) and structured illumination microscopy (SIM) to visualize cargo uptake. The isolated exosomes were also transfected with pre-miR-21 and filamin A (FlnA)-siRNAs before being co-cultured with podocytes. We confirmed the efficiency of transfection and knockdown using RT-qPCR, Western blotting, and immunofluorescence staining.

TEM revealed that the exosomes maintained a consistent shape and size of approximately 20 nm post-transfection and exhibited a stable expression of CD9 and TSG101. Our imaging showed that podocytes take up Cy3-labeled exosomal miRNAs and siRNAs, utilizing various mechanisms, including encapsulation within vesicular structures, endocytosis and free distribution within the cells. Transfection of exosomes with FlnA-siRNAs resulted in a significant 2.8-fold reduction of FlnA expression in co-cultured podocytes, while pre-miR-21-transfected exosomes led to a remarkable 338-fold increase in mature miR-21 levels.

These findings demonstrate that direct exosome transfection with fluorescently labeled small RNAs is an effective tool for exosomal cargo tracking in podocytes. This study is the first to show that directly transfected exosomes can deliver small RNAs to podocytes in vitro, suggesting their potential as RNA carriers for therapeutic strategies in more complex settings.

## 1. Introduction

Podocytes, a highly-specialized renal epithelial cell type, cover the outer aspect of the glomerular capillaries and are crucial for the size-selectivity of the filtration barrier. Changes of the complex 3D morphology of their interdigitating foot processes or their loss are the leading cause for 80% of chronic kidney diseases (CKD) (1).

CKD displays a major public health burden affecting more than 10% of the world’s population (2). Until now there is no causal therapy for the treatment of CKD patients (2). If CKD progresses, renal replacement therapies such as dialysis and transplantation are necessary. However, they are often associated with significant limitations in quality of life and with a higher mortality. Therefore, there is an urgent need for advanced strategies to generate new treatment options.

The study of extracellular vesicles, in particular exosomes, is an innovative and promising approach for the treatment of CKD. These vesicles play an important role in cell-cell communication between podocytes and could also act as natural carriers for a variety of molecular cargos – such as proteins, metabolites, DNA, RNA and small RNAs (3). Since it was shown that podocytes release increased amounts of exosomes containing disease-specific small RNA cargo during kidney injury (4), especially miRNAs recently came into the focus of kidney research (5, 6). Furthermore, there is growing evidence suggesting that exosomes could also function as delivery vehicles for small RNAs to podocytes. This would offer promising avenues for therapeutic interventions (7).

Current studies primarily use indirect loading strategies, such as exosomes isolated from naturally overexpressing cell types like stem cells, to treat podocyte-related diseases (8). Another common approach involves transfection of donor cells with specific small RNAs and the subsequent isolation of the enriched exosomes which they secrete (9). Both methods have major disadvantages: they lack specificity due to the presence of other endogenous small RNAs, often have insufficient intracellular packaging of the target small RNAs, reduced reproducibility, and the transfection reagents used are hardly removable and often toxic to recipient cells (10).

A more promising approach than the currently preferred indirect methods could be the direct transfection of exosomes with small RNAs. However, it is not yet known whether cultured podocytes can effectively internalize these cargo-loaded exosomes and whether the contained small RNAs remain functional. To address these questions, we directly transfected exosomes with fluorescently labeled miRNAs and siRNAs, demonstrating their potential as a viable tool for delivering small RNAs into podocytes.

## 2. Material and methods

### 2.1 Cell culture

Conditionally immortalized podocytes (SVI; CLS Cell Line Service GmbH, Eppelheim, Germany) were used for all cell culture experiments and have been handled as previously described (11). Podocytes were maintained in RPMI 1640 medium (Sigma-Aldrich, St. Louis, MO, USA) supplemented with 10% fetal bovine serum (FBS, Boehringer Mannheim, Mannheim, Germany), 100 U/mL penicillin, and 0.1 mg/mL streptomycin (Thermo Fisher Scientific, Waltham, MA, USA). For expansion, podocytes were cultured at 33°C and 5% CO_2_. To induce podocyte differentiation, podocytes were cultured at 38°C and 5% CO_2_ for at least two weeks before the experiments. Prior to exosome isolation, cells were washed three times with Dulbeccòs PBS (PBS, Sigma-Aldrich) and medium was replaced with RPMI 1640, supplemented with 10 % exosome-depleted FBS (System Biosciences, Palo Alto, CA, USA), 100 U/mL penicillin and 0.1 mg/mL streptomycin.

### 2.2 Exosome isolation

Cells were kept in exosome-depleted media for 3 days. The conditioned medium was transferred to a 15 mL conical tube and centrifuged for 15 min at 3000 x *g* at room temperature (RT) to remove cells and cell debris. The supernatant was transferred to a fresh 15 mL conical tube and was centrifuged again for 15 min at 3000 x *g* at RT to remove residual cell debris. Afterwards 2 mL Exoquick TC (System Biosciences) were added to the supernatant. After inverting the tube several times, it was incubated at 4°C overnight. Afterwards, the supernatant has been centrifuged for 1 hour at 10000 x *g* at 4°C. The exosome pellet was eluted in RPMI 1640 medium without supplements.

### 2.3 Exosome transfection

Exosome transfection was performed with the Exo-Fect™ siRNA/miRNA Transfection Kit (System Biosciences) after manufacturer’s instructions with some modifications. We used the Silencer Select Flna siRNA (#s101260, Ambion, Thermo Fisher Scientific), pre-miR™ miRNA Precursor miR-21 mimics (#AM17100, Ambion, Thermo Fisher Scientific) and the corresponding Cy3-labeled negative controls (Silencer™ Cy3™-labeled Negative Control No. 1 siRNA and Cy3™-labeled Pre-miR Negative Control #1, Ambion, Thermo Fisher Scientific). The volume of the transfection reaction was 110 µL and included 545 nM of each nucleic acid. After incubation for 15 min at RT, the transfection reaction mixture was applied to 100 µL eluted exosomes or PBS as a negative control, respectively. This mixture was incubated for 1 h at 37°C in the dark. Exosome cleanup was performed after manufacturer’s instructions.

### 2.4 Exosome treatment

Prior to exosome treatments, differentiated podocytes were washed twice with 10 mL PBS and cells were dissociated by 3 mL trypsin-EDTA. The trypsin-EDTA reaction was blocked by addition of 9 mL RPMI 1640 supplemented with exosome-depleted FBS, 100 U/mL penicillin and 0.1 mg/mL streptomycin. Subsequently, the cells were pelleted by centrifugation at RT and 750 x *g* for 5 min. The pellet was eluted in 1 mL RPMI 1640 supplemented with exosome-depleted FBS, 100 U/mL penicillin and 0.1 mg/mL streptomycin. After cell counting, 2.5×10^4^ cells per well were seeded in each well of a collagen 4-coated (Thermo Fisher Scientific) 6-well plate in 2 mL RPMI 1640, supplemented with exosome-depleted FBS, 100 U/mL penicillin and 0.1 mg/mL streptomycin. The cells were used for exosome treatments after 3 days. The cleaned-up exosome elutions from the transfection experiments were applied to the corresponding wells for different time spans.

### 2.5 Protein isolation and Western blot

Protein isolation of cells was performed as previously described (12, 13). Prior to protein isolation, cells were washed 3 times with 2 mL PBS.

For protein isolation of transfected and untransfected exosomes 100 µL of exosome elution were supplemented with 20 µL Exoquick TC, inverted and incubated on ice for 1 h. Afterwards samples were centrifuged for 1 h at 14000 x *g* and 4°C. The pellets were eluted in Pierce IP Lysis Buffer (500 mM, Thermo Fisher Scientific). Further protein isolation and concentration determination via Bradford assay were performed as previously described (13, 12).

Then, the samples were adjusted to 20 μg/lane (TSG101) and to 10 μg/lane (CD9) for qualitative analysis. For quantitative analysis we used 20 µL of each exosome sample per lane. For the analysis of cultured podocytes after treatment with FlnA-siRNA exosomes, we adjusted all samples to 2 µg per lane. All samples were mixed with 6 x sample buffer (0.35 M Tris [pH 6.8], 0.35 M SDS, 30% v/v Glycerol, 0.175 mM bromophenol blue) and boiled at 95°C for 5 min. The protein samples were separated on a 4–20% gradient Mini-Protean TGX Gel Stain-free (Bio-Rad, Hercules, CA, USA). The separated proteins were blotted on nitrocellulose membranes using the Trans-Blot Turbo RTA Transfer Kit (Bio-Rad) and the Trans-Blot Turbo Transfer System (Bio-Rad) at 2.5 A/25 V for 5 min. Membranes were washed in 1x TBS+T wash buffer (50 mM Tris, 150 mM NaCl, 10 mM CaCl_2_, 1 mM MgCl_2_ supplemented with 0.1% Tween-20; AppliChem, Darmstadt, Germany) and blocked in wash buffer supplemented with 5% milk powder (blocking solution) for 1 hour at room temperature. Primary antibodies were diluted in blocking solution and incubated with the membranes overnight. After washing 3 x 5 min with wash buffer, the membranes were incubated with secondary antibodies for 45 min, washed again 4 x 5 min with wash buffer, developed with the ECL Prime Western Blotting Detection Reagent (Cytiva Europe GmbH, Freiburg, Germany) and visualized on X-ray films (Cytiva Europe GmbH) by using Carestream Kodak autoradiography GBX developer/fixer solutions. For normalization and usage of alternative antibodies on the same blot, blots were stripped. The following antibodies were used at the final concentrations: anti-TSG101 (Sigma-Aldrich; 1:1000), anti-CD9 (Thermo Fisher Scientific, 1:2000), anti-FlnA (Sigma-Aldrich; 1:4000), anti-Gapdh (Santa Cruz Biotechnology, Santa Cruz, USA; 1:2000), secondary anti-rabbit HRP (Santa Cruz Biotechnology; 1:6000 for TSG101; 1:5000 for CD9; 1:15000 for FlnA and Gapdh).

### 2.6 Transmission electron microscopy

For transmission electron microscopy (TEM), 10 mL of cell- and debris-free cell culture media were treated with ExoQuick-TC (System Biosciences) for exosome isolation purposes according to the manufacturer’s instructions with minor modifications. To 10 mL cell culture media, 3.3 mL ExoQuick-TC were added and stored overnight at 4°C. Then, samples were centrifuged at 10000 x *g* for 60 min and the supernatant was discarded. The pellet was prepared for TEM according to Asadi and coworkers (14) using the flotation method for the staining procedure. Briefly, isolated exosomes were fixed with 2% paraformaldehyde in 0.1 M sodium phosphate buffer (pH 7.5) and then allowed to adsorb onto a glow-discharged carbon-coated holey Pioloform film on a 400-mesh grid (Plano GmbH) for 20 min on ice. The grid was then transferred onto four droplets of deionized water on ice for 2 min each, and finally onto a drop of staining mixture for 10 min on ice. To prepare the staining mixture, 100 µL of 3% aqueous uranyl acetate were added to 900 µL 2% methylcellulose followed by the addition of 75 µL deionized water and 25 µL 1% phosphotungstic acid (pH 7). After blotting with filter paper, the grids were air-dried. All samples were examined with a transmission electron microscope LEO 906 (Carl Zeiss Microscopy Deutschland GmbH, Oberkochen, Germany) at an acceleration voltage of 80 kV. All micrographs were edited by using Adobe Photoshop CS6.

### 2.7 RNA isolation

For miR-loading experiments cells were cultured for 24 h in the presence of transfected and untransfected exosomes and controls as indicated. For siRNA-loading experiments cells were treated for 48 h with transfected exosomes and controls as stated. RNA isolation was performed as previously described (12). Prior to administration of Tri-reagent the cell layer was washed twice with 2 mL PBS, respectively.

### 2.8 Taqman reverse transcription

cDNA synthesis was performed starting with 10 ng total RNA using Taqman™ miRNA Assays and the Taqman™ miRNA Reverse Transcription Kit (Thermo Fisher Scientific). The following Taqman™ miRNA Assays were used: Hsa-miR-21-5p: ID #000397 and U6 snRNA: ID #715680. The RT-reactions were performed after manufacturer’s instructions. Negative controls included no template and no reverse transcriptase controls.

### 2.9 Taqman qPCR

The qPCR was performed with the Taqman™ miRNA Assays mentioned above and the Taqman™ Universal Master Mix II, no UNG (Thermo Fisher Scientific) following the manufacturer’s instructions. The reaction volumes contained 1.33 μL undiluted cDNA solution and 18.67 μL Master Mix. The qPCR was performed on a Bio-Rad iQ5 thermal cycler (Bio-Rad) with the following cycler scheme: 10 min at 95°C followed by 45 cycles of 15 sec at 95°C and 60 sec at 60°C. All samples were run in triplicate.

Negative controls included the no template controls from cDNA synthesis and an extra no template control for the qPCR reaction. Ct-values were calculated with automatically set thresholds and baselines by the cycler software (Bio-Rad). Raw Ct-values ≥ 38 were excluded from analysis. All Ct-values were normalized against U6 and the Cy3 w/o exosomes Ctrl.

### 2.10 Immunofluorescence staining

Prior to immunofluorescence staining cells were fixed with 2% paraformaldehyde (PFA) for 10 min and were permeabilized with 0.3% Triton-X (Sigma-Aldrich) for 3 min. Then cells were blocked for 1 hour with blocking solution (2% FBS, 2% BSA and 0.2% fish gelatin in PBS). Primary antibodies were diluted 1:100 in blocking solution and were incubated for 1 hour at RT. We used the following primary antibodies: anti-FlnA (Sigma-Aldrich), anti-CD9 (BD Biosciences, San Jose, CA), anti-Rab5 (Cell Signaling Technologies, Leiden, Netherlands) and anti-Rab7 (Cell Signaling Technologies). Secondary antibodies were diluted 1:300 in blocking solution and were incubated with the cells for 1 h: anti-mouse-Cy2 (Jackson Immuno Research Laboratories, Ely, UK) and anti-rabbit-Cy2 (Jackson Immuno Research Laboratories). The actin cytoskeleton was visualized by staining with Alexa Fluor 647 phalloidin (1:100; Thermo Fisher Scientific) for 1 h that was added to the secondary antibody solution. For nuclei staining DAPI diluted 1:100 in PBS (Sigma-Aldrich) was used for 2 min. All samples were mounted in Mowiol (Carl Roth, Karlsruhe, Germany).

### 2.11 Confocal laser scanning microscopy (cLSM)

Confocal laser scanning microscopy (cLSM) was performed on a Leica TCS SP5 confocal microscope (Leica Microsystems, Wetzlar, Germany) with 20x, 40x and 63x oil immersion objectives. For image acquisition the Leica Application Suite software (Leica Microsystems, Version 2.6.0) was used. All z-stacks were acquired throughout the whole z-plane (20 µm) in 0.5 µm steps. Pre-miR-21 experiments were acquired in 20 x 1µm steps. The xzy-projection was acquired in 20 x 1.73 µm steps. For 3D-SIM a Zeiss Elyra PS.1 System (Zeiss Microscopy, Oberkochen, Germany) equipped with a 63x oil immersion objective was used. Z-Stacks with a size of 1280 µm × 1280 µm with a slice-to-slice distance of 0.43 µm were acquired over approximately 2 µm using the 561 nm laser, (3.5% laser power, exposure time: 150 ms), the 488 nm laser (3.5% laser power, exposure time: 150 ms) and the 405 nm laser (7% laser power, exposure time: 100 ms). The grating was shifted and rotated five times on every frame while acquiring widefield images. Z-stacks were acquired in 4 planes with 0.43 µm. 3D-SIM reconstruction was performed with the Zeiss ZEN Black. ZEN Blue software was used for maximum intensity projections.

### 2.11 Statistical analysis

Statistical analysis was performed using GraphPad prism V5.01 (GraphPad Software, CA, USA, https://www.graphpad.com). Data was checked for gaussian distribution by Kolmogorov–Smirnov test. All groups were tested for statistically significant differences by two-way ANOVA and Bonferroni post-hoc test. All values are displayed as means with standard deviations. P-values□≤□0.05 were considered as statistically significant.

## 3. Results

### 3.1 Transfected exosomes show a typical morphology

To characterize exosomes derived from cell culture supernatants before and after transfection with the Exo-Fect™ siRNA/miRNA Transfection Kit (System Biosciences), exosome pellets were prepared for TEM analysis. We observed that the exosomes exhibit the characteristically cup-shaped morphology with a size ranging between 20 - 30 nm. Interestingly, exosomes showed a tendency to aggregate and to form clusters before transfection, a phenomenon absent in transfected exosomes. These exosomes displayed a homogenous distribution and were distinct from another. However, we did not observe differences in exosome number or size (Fig. 1 A).

**Figure 1:**
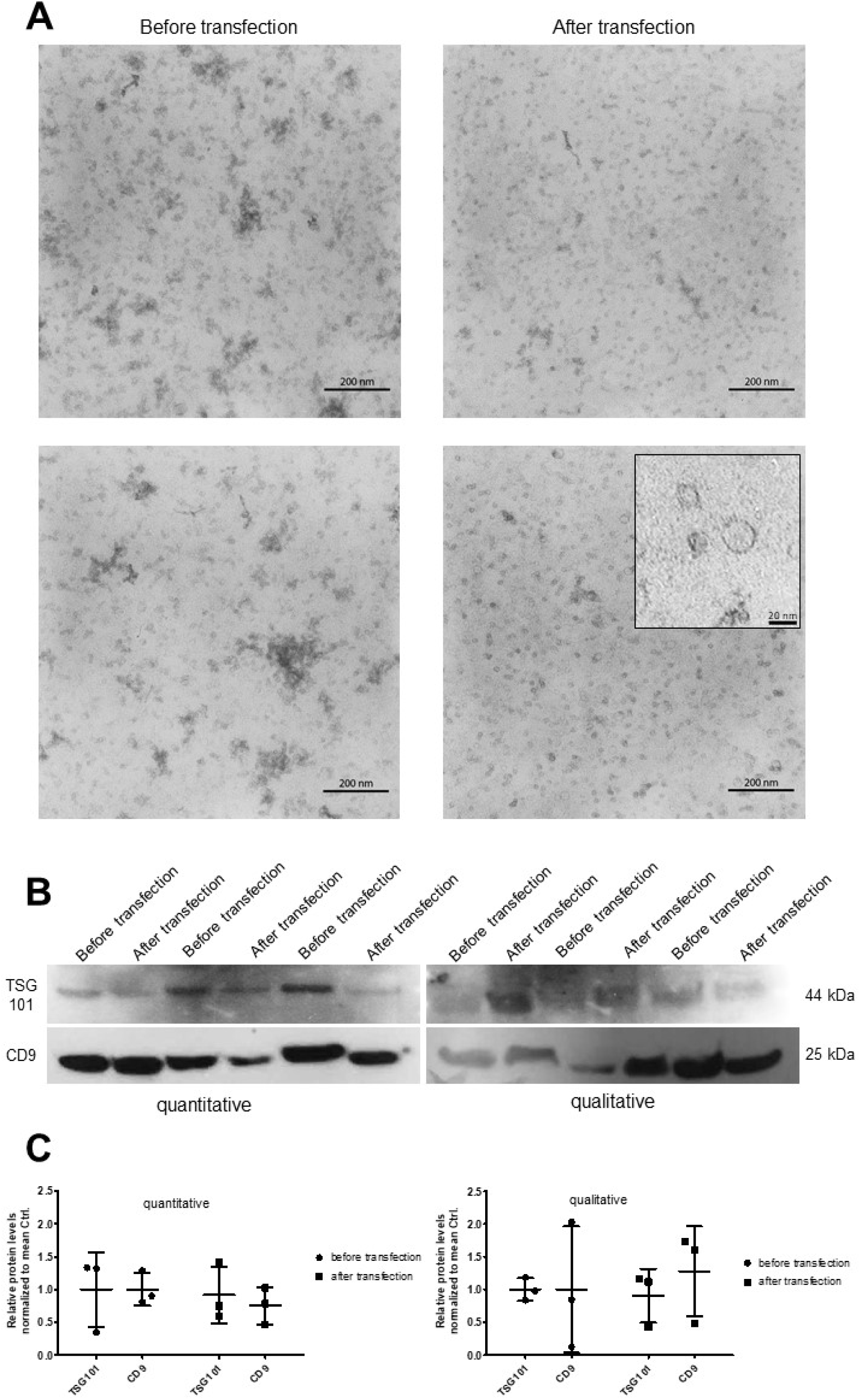
Exosome characterization before and after transfection. (A) Exosomes were isolated with ExoQuick-TC before and after transfection with Cy3-miRCtrl with the Exo-Fect™ siRNA/miRNA Transfection Kit. Exosomes were prepared for TEM. Isolated exosomes showed a typical shape and size of 20-30 nm and shape (box). There was no difference in exosome number after transfection. Exosomes appeared cleaner and more homogeneously distributed after transfection. Scale bar = 200 nm, box = 20 nm. (B) Exosomes of three independent transfections with Cy3-miRCtrl were precipitated with ExoQuick-TC and analyzed by Western blot for exosomal marker proteins TSG101 and CD9 quantitatively and qualitatively. There are no differences in exosomal marker levels in both analysis approaches. (C) The bands of the Western blots were analyzed semi-quantitatively by using Fiji. Data shown are means ± SD. There were no significant differences in exosomal marker expression before and after transfection of exosomes.

### 3.2 Transfected exosomes show a typical marker expression

To further characterize isolated exosomes, we performed qualitative as well as quantitative Western blots for the specific exosomal marker proteins TSG101 and CD9 (Fig. 1 B, C). The quantitative Western blots (equal volumes) showed a slight, non-significant reduction of TSG101 and CD9 expression of transfected exosomes (Fig. 1 B, C) indicating a reduced exosome amount after transfection. Regarding the qualitative (equal protein input) Western blots, we detected slightly increased CD9 signals in exosomes post transfection compared to exosomes prior to transfection, indicating a higher exosome purity after transfection.

### 3.3 Podocyte uptake of exosomes loaded with Cy3-labeled pre-miRNA

We incubated cultured murine podocytes with Cy3-labeled control miRNA-loaded exosomes (Cy3-miRCtrlExos) for 48 hours. To rule out any background staining from unloaded Cy3-fluorophores, we incubated the podocytes with Cy3-labeled Ctrl-miRNA without exosomes (Cy3-miRCtrl w/o Exos) that had undergone the same washing procedure as transfected exosomes, followed by a phalloidin and DAPI staining. We observed an internalization of Cy3-miRCtrlExos by podocytes using cLSM (Fig. 2 A). This was additionally confirmed by cLSM in xzy-plane (Fig. 2 B). Since the Cy3-signals were found in close vicinity to F-actin filaments, we assumed that the exosomal miRNA cargo was taken up by the cultured podocytes. In contrast, Cy3-signals were rarely detected in samples treated only with Cy3-miRCtrl w/o Exos. These results indicate the selectivity of the exosome transfection procedure and suggest that unloaded Cy3-miRNA signals do not play a significant role in this context.

**Figure 2:**
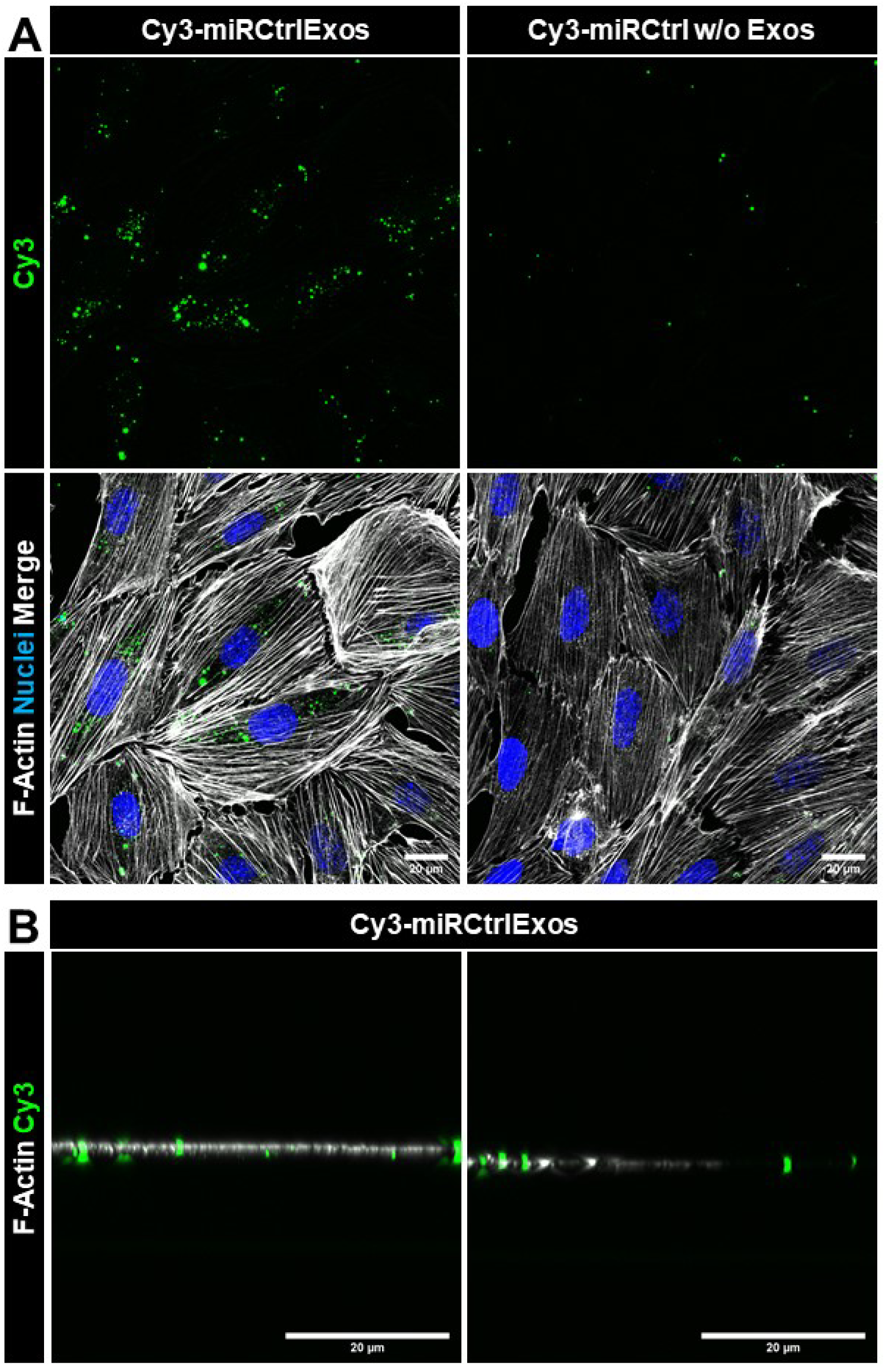
Cultured podocytes internalize cargo of exosomes loaded with Cy3-Ctrl miRNA. (A) Cultured murine podocytes were treated for 48 hours with exosomes previously transfected with Cy3-Ctrl-miRNA (Cy3-miRCtrlExos) and with Cy3-Ctrl-miRNA w/o exosomes (Cy3-miRCtrl w/o Exos). Podocytes treated with Cy3-miRCtrl w/o Exos showed only a few Cy3 spots. Podocytes treated with transfected exosomes showed intracellular Cy3-signals as shown by xyz (A) as well as by (B) xzy-projections. Scale bars = 20 µM. F-Actin is stained with phalloidin. Nuclei are stained with DAPI. Maximum intensity projections (MIP) of z-stacks.

### 3.4 The internalization of exosomal cargo is confirmed by co-staining against CD9, Rab5 and Rab7

To further confirm internalization of Cy3-miRCtrlExos, we stained podocytes with a specific antibody against the exosomal marker protein CD9. Hereby, we found that most of the Cy3-positive signals are localized within the podocyte cell bodies (Fig. 3 A). Interestingly, we detected CD9-negative Cy3-spots (Fig. 3 A, arrow) as well as Cy3-signals that were encapsulated in CD9-positive vesicular structures at higher magnifications (Fig. 3 A, asterisk, B). The diameter of these vesicular structures varied between 200 and 700 nm.

**Figure 3:**
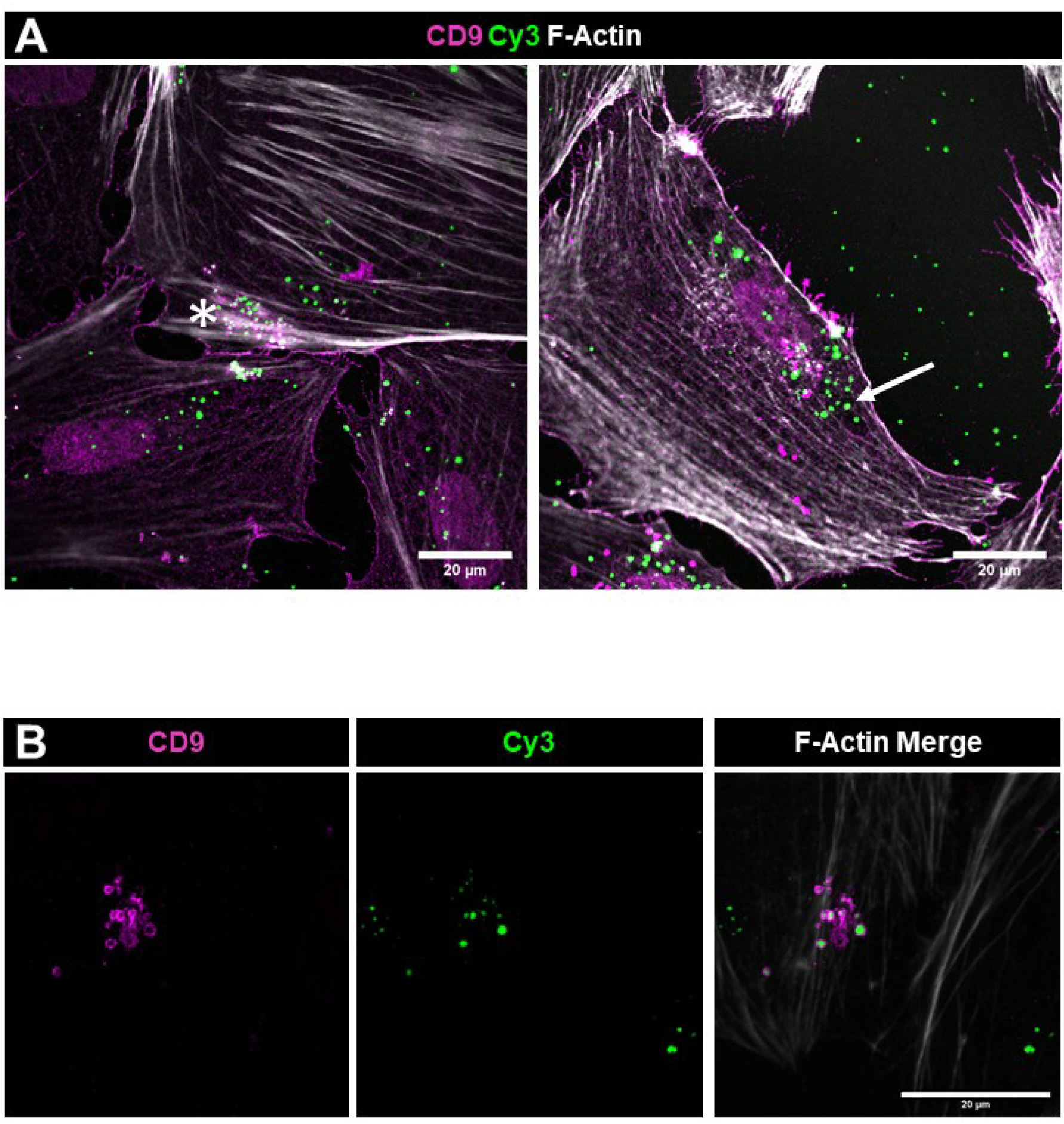
Staining for CD9 confirms the internalization of exosomal cargo. (A) After the treatment with Cy3-miRCtrlExos, podocytes were stained with an antibody against CD9. Podocytes showed Cy3-signals inside the cell body. The Cy3-signals were either CD9-negative (A, arrow) or enclosed in CD9-positive vesicles (A, asterisk and B). F-Actin is stained with phalloidin. Scale bars = 20 µM. Single planes (A) and MIP of z-stacks (B).

To further elucidate the mechanism of the exosomal cargo uptake, we stained our cells with an antibody against the early endosome marker Rab5. By using super resolution microscopy SIM, we observed a broad co-localization of Cy3 and Rab5 (Fig. 4 A). A staining against Rab7, a late-endosome and lysosome marker protein, showed a fewer co-localization with Cy3-signals compared to Rab5 (Fig. 4 B).

**Figure 4:**
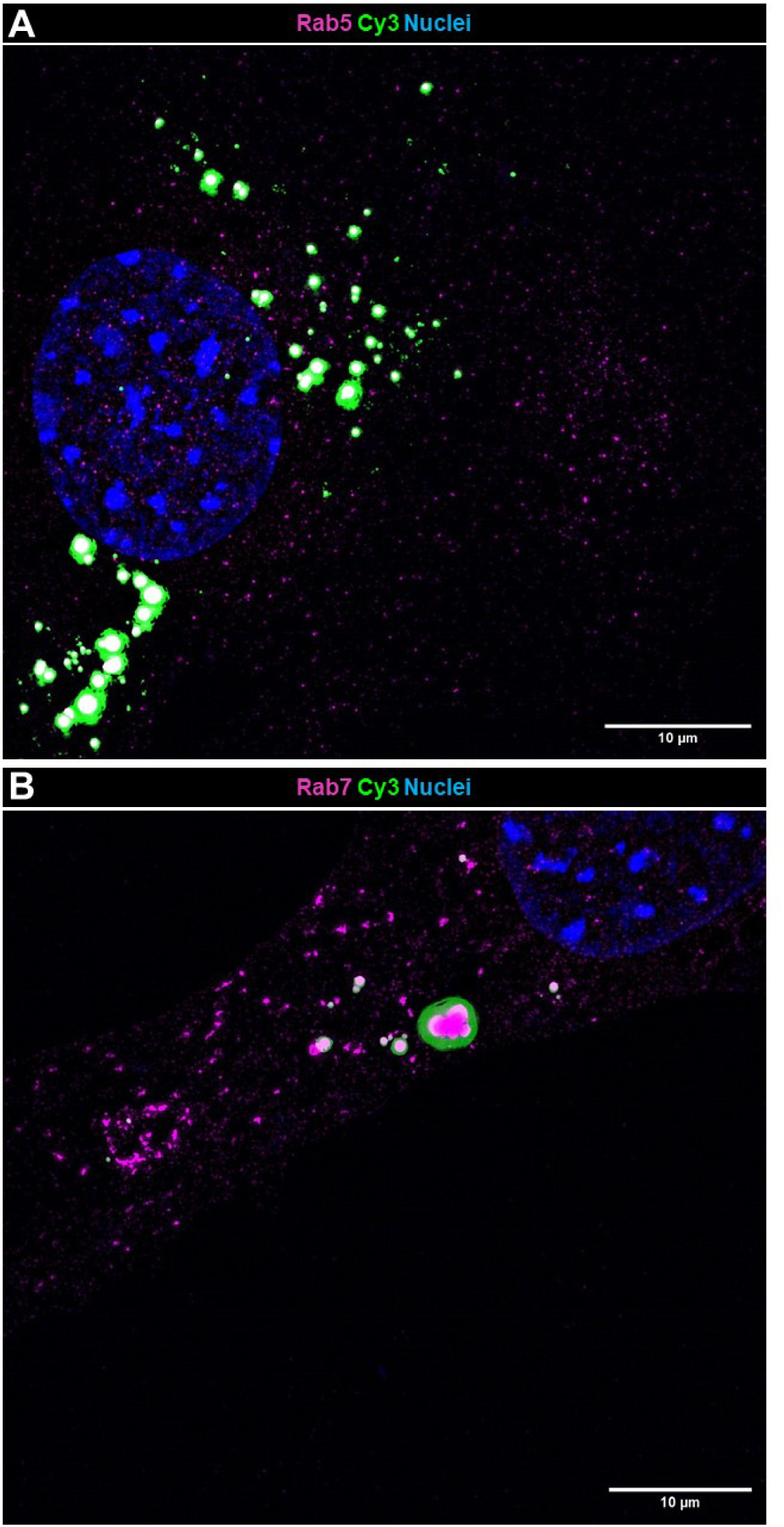
Staining for the early endosomal marker Rab5 and the lysomal marker Rab7 reveals endocytotic exosome cargo uptake rather than lysosomal degradation. Cultured podocytes were stained with antibodies against Rab5 (A) and Rab7 (B) after the treatment with Cy3-miRCtrlExos. (A) Podocytes showed a wide co-localization of Cy3 with the early endosomal marker Rab5. (B) Cy3-positive signals were rarely co-localized with the lysosomal marker Rab7. To visualize small particles, microscope settings were set to signal saturation. Nuclei are stained with DAPI. Scale bars = 20 µM. MIP of z-stacks.

### 3.5 Time dependence of the exosomal cargo internalization

To investigate the dynamics of exosomal cargo internalization, we treated podocytes with Cy3-miRCtrlExos starting from 24 hours up to 1 week. We observed Cy3-Ctrl signals internalized by the cells as well as in the cell-free periphery to equal amounts after 24 hours treatment (Fig. 5). We identified Cy3-miRCtrl signals extracellularly as well as internalized by the podocytes. Additionally, we observed an accumulation of Cy3-Ctrl signals in close proximity of cell borders as seen in higher magnification in Fig. 5. After 48 h, only a few Cy3-Ctrl signals were visible in the periphery whereas Cy3-Ctrl signals accumulated in the cellular area (Fig. 5). Over the period of one week, Cy3-Ctrl signals further disappeared from the cell-free areas, and all detected Cy3-Ctrl signals were localized within the cells and partially accumulated in CD9-positive vesicular structures (Fig. 5).

**Figure 5:**
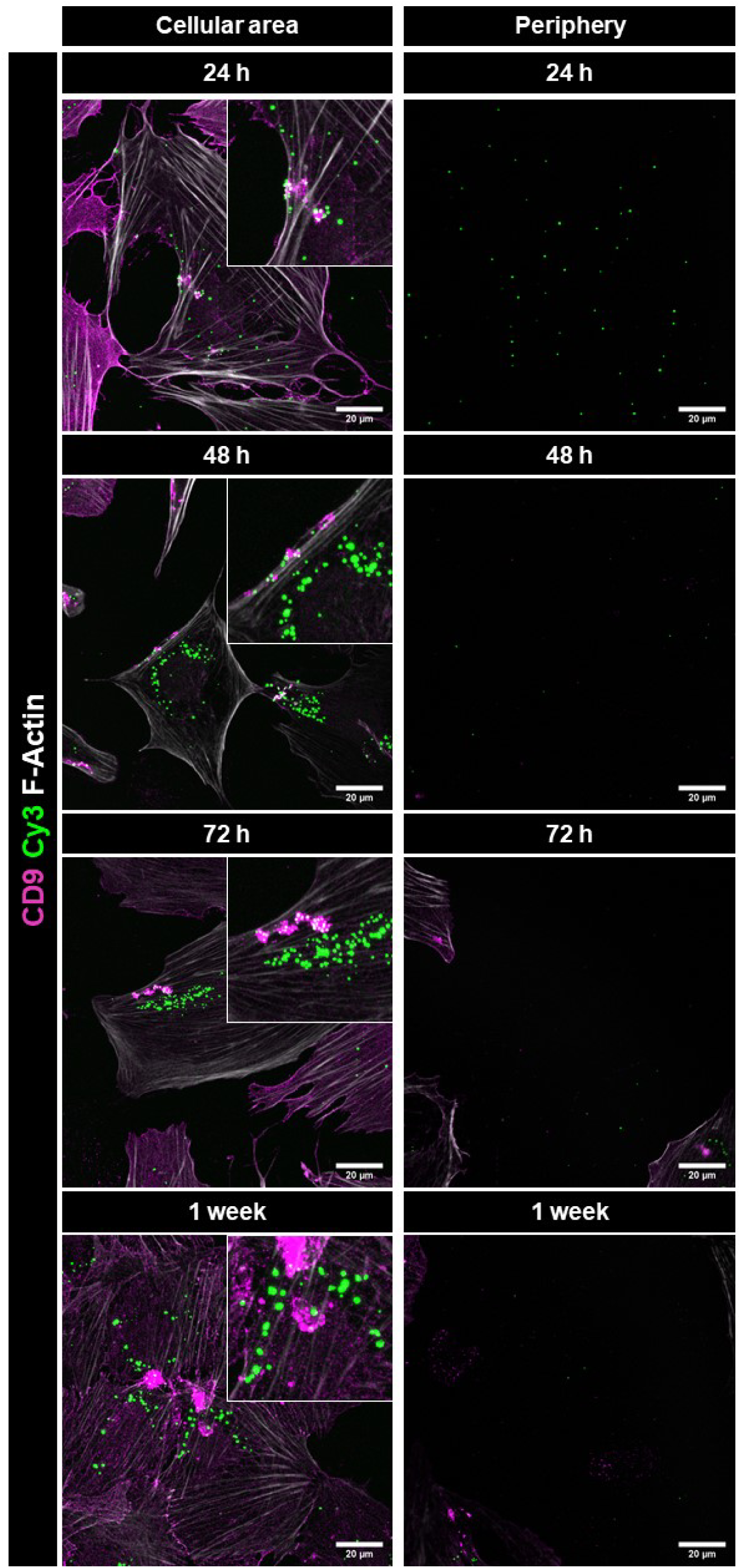
Time dependence of the exosomal cargo internalization. Podocytes were treated with Cy3-miRCtrlExos for 24, 48, 72 hours and 1 week followed by a staining against CD9. Immunofluorescence micrographs were acquired from the cellular area as well as from the cell-free periphery. The number of Cy3-spots decreased in the periphery and increased within the cells over time. F-Actin is stained with phalloidin. Scale bars = 20 µM. MIP of z-stacks (cellular area) and single plane (periphery).

### 3.6 Treatment of podocytes with pre-miR-21-loaded exosomes leads to overexpression of mature miR-21

After successful exosomal cargo tracking in podocytes, we investigated whether exosomes could serve as delivery vehicles for pre-miRNAs. Therefore, we treated cultured podocytes with exosomes transfected with pre-miR-21 (pre-miR-21Exos) and Cy3-miRCtrl (Cy3-miRCtrlExos). We analyzed the internalization efficiency by cLSM. To evaluate whether the loaded pre-miR-21 gets processed to its mature form we isolated total RNA from treated podocytes and measured the amount of mature miR-21 by RT-qPCR. Prior to this, we performed several washing steps to eliminate uninternalized exosomes. We found Cy3-signals within all podocytes treated with Cy3-miRCtrlExos indicating full internalization of the exosomal cargo (Fig. 6 A). Podocytes treated with unlabeled pre-miR-21Exos were free of Cy3 fluorescence. By RT-qPCR, we found a significant 338-fold (p = 0.03) up-regulation of mature miR-21 compared to Cy3-miRCtrl w/o Exos-treated podocytes. In contrast, podocytes treated with Cy3-miRCtrlExos showed no significant difference in mature miR-21 expression levels (0.71-fold, p = 0.15) compared to Cy3-miRCtrl w/o Exos Ctrl. However, they differed significantly from pre-miR-21Exos-treated samples (p = 0.03) (Fig. 6 B).

**Figure 6:**
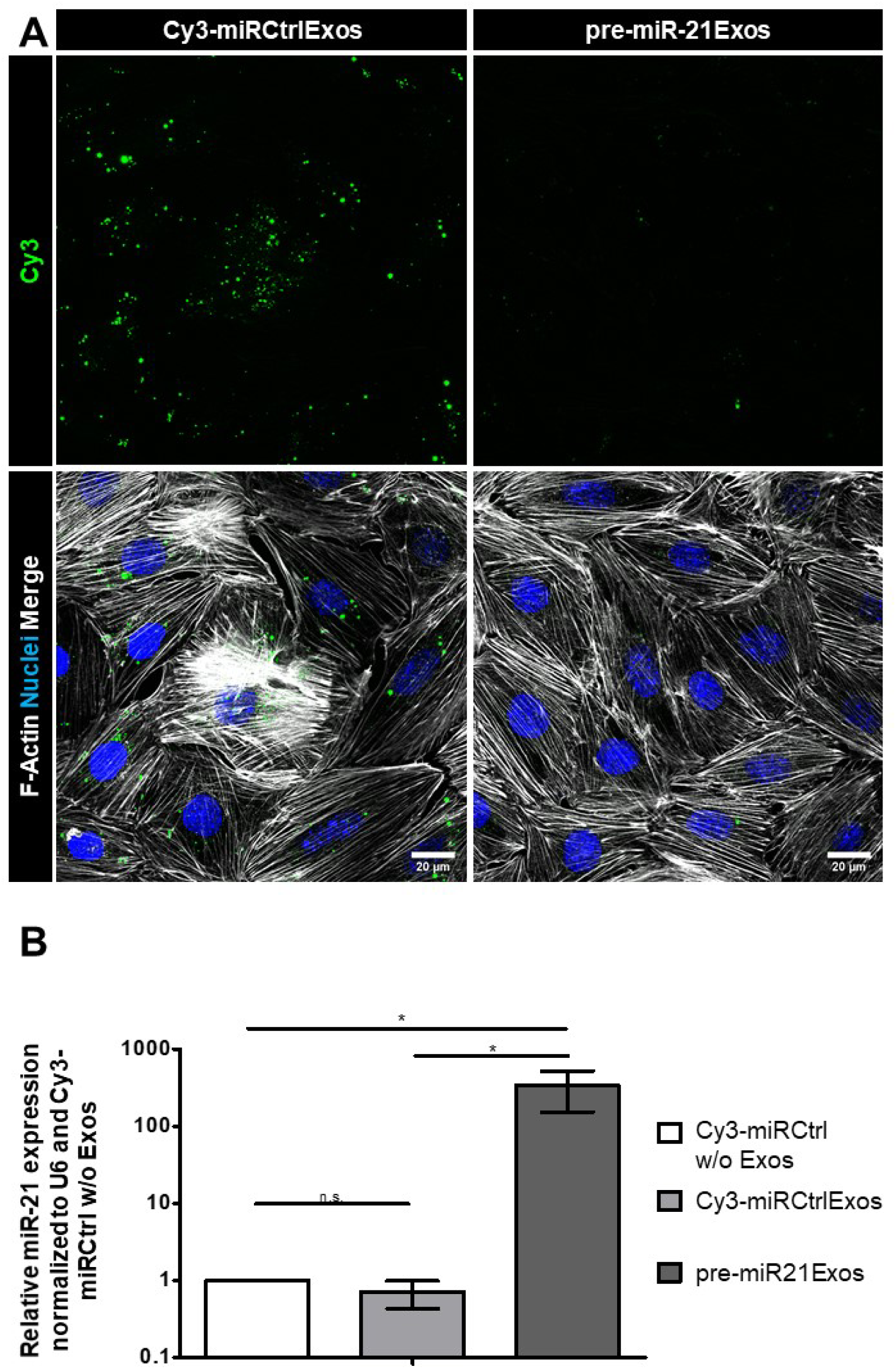
Exosomes loaded with pre-miR-21 up-regulate miR-21 expression in podocytes. (A) Podocytes were treated with Cy3-miRCtrlExos and pre-miR-21Exos for 48 hours. Podocytes internalized Cy3-Ctrl, whereas almost no Cy3-fluorescence was detected in unlabeled pre-miR-21Exos-treated samples. F-Actin is stained with phalloidin. Nuclei are stained with DAPI. Scale bars = 20 µM. MIP of z-stacks. (B) Treatment with pre-miR-21Exos led to a 338-fold up-regulation of miR-21 in podocytes (p = 0.03). Graph shows the relative miR-21 expression normalized to U6 and Cy3-miRCtrl w/o Exos as mean ± SD.

### 3.7 Treatment of podocytes with Filamin A siRNA-loaded exosomes leads to the down-regulation of Filamin A protein expression

Besides pre-miRs as cargo molecules, we investigated if siRNAs could also be successfully transferred to cultured podocytes by exosome transfection. For this purpose, we used a well-established siRNA against Filamin A (FlnA) mRNA as a representative target (FlnA-siRNAExos). Furthermore, we used Cy3-labeled, scrambled siRNA-Ctrl as an internalization control (Cy3-siRNACtrlExos). In the comparative analysis shown in Figure 7 A, podocytes treated with Cy3-siRNACtrlExos exhibited fewer and weaker Cy3 signals than cells treated with Cy3-miRCtrlExos (Fig. 6A). Remarkably, podocytes treated with FlnA-siRNAExos showed a complete absence of Cy3 signals which looks very similar to the untreated control. (Fig. 7 A). This finding indicates a very reliable exosome clean-up. After the incubation with FlnA-siRNAExos, we found a down-regulation of FlnA in contrast to Cy3-siRNACtrlExos-treated and untreated cells (Fig. 7 A), where no change of the FlnA expression was detected. Approximately 50 % of the treated podocytes per field of view showed almost no FlnA protein staining, whereas all control cells were FlnA-positive. These results were confirmed by Western blot analysis which showed a significant (p = 0.006) decrease to 0.36 in FlnA expression in the samples treated with FlnA-siRNAExos, compared with Cy3-siRNACtrlExos and untreated controls (Fig. 7 B).

**Figure 7:**
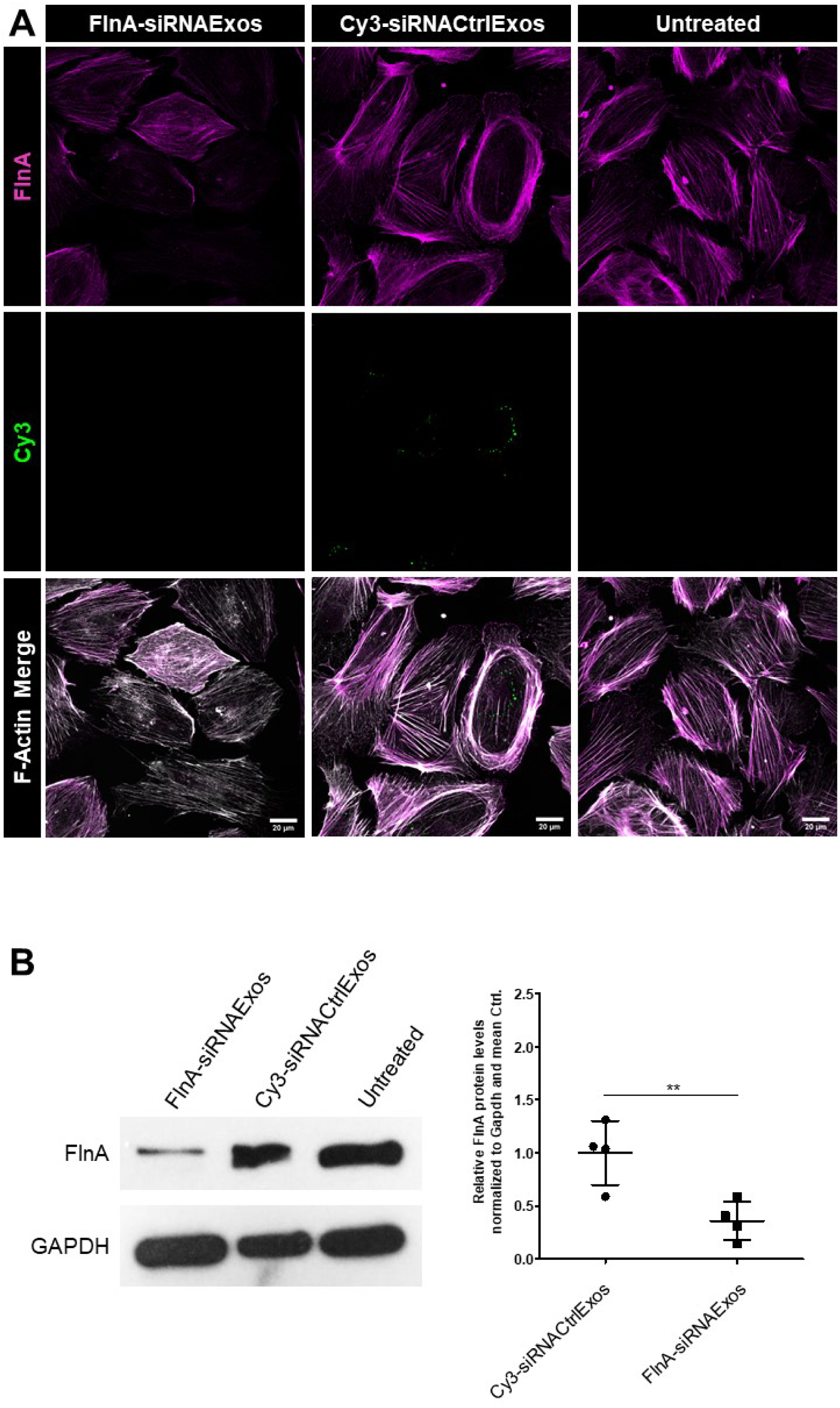
Exosomes loaded with FlnA-siRNA down-regulate FlnA expression in podocytes. (A) Podocytes were treated with Cy3-siRNACtrlExos and FlnA-siRNAExos for 48 hours. Podocytes internalized Cy3-siRNACtrl while no Cy3 signals were found in untreated and FlnA-siRNAExos samples. FlnA fluorescence decreased in FlnA-siRNAExos-treated samples. F-Actin was stained with phalloidin. Scale bars = 20 µm. MIP of z-stacks. (B) Treatment of podocytes with FlnA-siRNAExos resulted in a down-regulation of FlnA expression to 0.36 (p = 0.006), shown by Western blot analysis.

## 4. Discussion

In the past, exosomes have been shown to be effective carriers for the transfer of small RNAs to different cell types, making them a promising therapeutic tool for the treatment of various diseases (7). However, in podocyte diseases, which account for 80% of all CKD cases, there are no treatment options based on specific target RNAs loaded into exosomes so far. However, this new approach could potentially close the large gap in the currently limited treatment options.

The present exosome-based studies mostly used indirect loading strategies. This includes exosomes isolated from special cell types that naturally overexpress RNAs, such as stem cells. In their study, Jin et al. treated diabetic mice with miR-486-5p-enriched exosomes secreted by adipose-derived stem cells, which resulted in an improvement of podocyte damage by regulating the Smad1/mTOR signaling pathway (8). Another common indirect approach is the transfection of donor cells with specific small RNAs and the subsequent isolation of enriched exosomes secreted by these cells. Zhang et al. transfected HEK cells with miR-145-5p, isolated miR-145-5p-enriched exosomes and then co-cultured these exosomes with podocytes. Thereby they observed toxic effects of the miR-145-5p-enriched vesicles on cultured podocytes (9, 8). However, both strategies have considerable disadvantages: Firstly, they lack specificity, as many other endogenous exosomal small RNAs are present in addition to the target RNAs. Secondly, the efficiency of packaging transfected small RNAs into exosomes within donor cells is often insufficient, resulting in lower transfection rates and poor reproducibility. Thirdly, transfection reagents are difficult to remove completely from the isolated exosomes and can therefore be cytotoxic to recipient cells (10).

A promising alternative approach is the direct transfection of exosomes, already being used in cancer research (15). However, it is still unclear whether directly transfected exosomes can effectively deliver small RNAs into podocytes. The aim of this study was to directly load exosomes isolated from cultured mouse podocytes with small RNAs which were then incubated with podocytes.

The isolated exosomes presented the classical cup-shape and size of 20-30 nm before and after transfection. This relatively small exosome size is typical for podocyte-derived exosomes and has been published previously (16, 17). Furthermore, there was no significant loss in the quantity or quality of exosome following transfection of the prepared exosomes. Interestingly, we noted a more homogenous exosome distribution after transfection via TEM.

To evaluate the efficiency of this method, we used fluorescently labeled miRNAs and siRNAs to track the uptake of exosomal cargo by the podocytes. Currently, most studies use staining protocols to visualize exosome uptake, by using fluorescent labeling of exosomal membrane components either via immunofluorescence or organic dyes (18). These methods have various disadvantages since organic dyes are predisposed to stain cellular components other than exosomes and thereby generate false positive results. Immunofluorescence staining using specific antibodies for exosome membrane proteins is very cost and labor intensive and includes various washing steps which drastically reduces exosome yield (18). In the present study, we circumvented these drawbacks by using prelabeled exosomal cargo instead of staining exosomal structural components. This technique not only increases the potential for tracking small RNA transport, but also offers significant advantages for tracking exosomes and exosomal cargo. By using this method, we were able to show that cultured podocytes internalize the fluorescently labeled small RNAs delivered by directly transfected exosomes.

To reveal the mechanism of uptake, we performed additional experiments. In general, exosomes can be internalized by different ways: by various forms of endocytosis – like clathrin-dependent (19, 20) and caveolae-dependent (21, 20) endocytosis (22, 20) and macro- and pinocytosis (19, 20) or receptor-mediated endocytosis. We have observed that the internalized small RNAs are localized either within the cytosol or encapsuled in CD9-positive vesicles. To determine the dominant vesicle type, we stained for Rab5, a marker for early endosomes (23) and Rab7 for late endosomes (24). After staining of podocytes with the specific antibodies, we found out that most Cy3 fluorescence colocalized with Rab5 and to a lesser extent with Rab7. This suggests that the uptake of exosomal cargo by podocytes *in vitro* occurs by endocytosis. However, the exact mechanisms of exosomal cargo uptake of podocytes remain rather unclear and require further studies.

To proof if transfected small RNAs remain functional, exosomes were loaded with specific siRNAs against FlnA as a representative target. FlnA has been reported to be essential for podocyte cytoskeleton integrity and is a good readout for RNAi efficiency since it is highly expressed in cultured murine podocytes (25). We found that podocytes which were treated with FlnA-siRNA exosomes, down-regulated FlnA. This verified that transfected siRNAs are functionally intact.

Likewise, a similar result was also observed for pre-miR-21-loaded exosomes. MiR-21 has been shown to be up-regulated in kidney injury and also in urinary exosomes of CKD patients (13). We found that mature miR-21 was strongly up-regulated in podocytes after pre-miR-21-exosome treatment.

During biogenesis of miRNAs, they undergo several processing steps. After transcription, miRNAs are present as pri-miRs with a length of > 1 kb and are further processed intranuclearly into 60 - 70 nt long pre-miRs with a stem-loop structure. These pre-miRs are exported into the cytosol and then cleaved by Dicers into mature miRNA and an opposing strand, which is further degraded (26). Using Taqman probes that are specific for mature miR-21 and can distinguish between pre-miRs and their mature forms with 2000-fold higher efficiency (27), we observed that the transfected pre-miR-21 was processed into mature miR-21 by the recipient podocytes, confirming their integrity. These results are consistent with previous findings that RNAs packed in exosomes are protected from degradation by RNases (28).

Our findings may open up great possibilities for various experimental approaches in podocyte research. The uptake of exosomes by podocytes has been reported several times before (29, 30) but to the best of our knowledge, this is the first time that one describes the small RNA uptake in podocytes by directly transfected exosomes.

Podocytes, a postmitotic cell type, undergo cell cycle arrest even under most *in vitro* conditions (31). Similar to other postmitotic cells, podocytes are difficult to transfect and highly sensitive to transfection reagents (31). Here we show that directly transfected exosomes could display a promising and gentle alternative to overcome these challenges. Direct exosome transfection could also play a potential, promising role for the targeted, therapeutic administration of small RNAs to podocytes in higher organisms. Evidence from the oncology research suggests that cells are more prone to phagocytize exosomes then liposomes for example, which are currently the gold standard in RNA delivery trials (32). Moreover, exosomes are less immunogenic then liposomes and express damage-specific surface markers that facilitate a specific uptake by injured cells (33).

In summary, we have shown that direct exosome loading with fluorescently-labeled small RNAs is suitable for exosome tracking approaches in podocytes *in vitro*. This study is the first to demonstrate the potential of directly transfected exosomes for delivering small RNAs to podocytes *in vitro,* providing an initial indication that exosomes could serve as small RNA-carriers for therapeutic strategies in more complex experimental settings.

## Acknowledgement

This work was supported by a grant of the Federal Ministry of Education and Research (BMBF, #01GM1518B, *“STOP-FSGS”*) and a grant of the Bundesministerium für Wirtschaft und Klimaschutz (BMWi, #16KN077229, *“AlternaTier-vivoPod”*). This work was supported by the Südmeyer fund for kidney and vascular research (*“Südmeyer Stiftung für Nieren-und Gefäßforschung”*) and the Dr. Gerhard Büchtemann fund, Hamburg, Germany.

The authors thank Annette Meuche for excellent technical assistance regarding electron microscopy.

